# Antagonism between the dynein and Ndc80 complexes at kinetochores controls the stability of kinetochore-microtubule attachments during mitosis

**DOI:** 10.1101/254359

**Authors:** Mohammed A. Amin, Richard J. McKenney, Dileep Varma

## Abstract

Chromosome alignment and segregation during mitosis depends critically on kinetochore-microtubule (kMT) attachments that are mediated by the function of the molecular motor cytoplasmic dynein, and the kinetochore microtubule (MT) binding complex, Ndc80. The RZZ (Rod-ZW10-Zwilch) complex is central to this coordination as it has an important role in dynein recruitment and has recently been reported to have a key function in the regulation of stable kMT attachment formation in *C. elegans*. However, the mechanism by which kMT attachments are controlled by the coordinated function of these protein complexes to drive chromosome motility during early mitosis is still unclear. In this manuscript, we provide evidence to show that Ndc80 and dynein directly antagonize each other’s MT-binding. We also find that severe chromosome alignment defects induced by depletion of dynein, or the dynein adapter spindly, are rescued by codepletion of the RZZ component, Rod, in human cells. Interestingly, the rescue of chromosome alignments defects was independent of Rod function in activation of the spindle assembly checkpoint and was accompanied by a remarkable restoration of stable kMT attachments. Furthermore, rescue of chromosome alignment was critically dependent on the plus-end-directed motility of CENP-E, as cells codepleted of CENP-E along with Rod and dynein were unable to establish stable kMT attachments or align their chromosomes properly. Taken together, our findings support the idea that the dynein motor may control the function of the Ndc80 complex in stabilizing kMT attachments either directly by interfering with Ndc80-MT binding, and/or indirectly by modulating the Rod-mediated inhibition of Ndc80.

## Introduction

Faithful chromosome segregation during mitosis requires proper chromosome congression, which relies on multiple mechanisms that ultimately lead to chromosome bi-orientation, an arrangement where the chromosomes are connected to microtubules from both spindle poles (1). In the classical ‘search and capture’ model, chromosomes move toward the spindle equator as a result of their biorientation (2). After nuclear envelope breakdown (NEB), chromosomes have been known to congress by two main mechanisms: microtubule polymerization/depolymerization-based motion (3) and motor-dependent transport along microtubules achieved by the coordinated activities of Dynein, CENP-E and Chromokinesins (4-6). The peripheral chromosomes are first transported by dynein to a microtubule-dense region near the spindle pole, from where they move towards the spindle equator along pre-existing spindle microtubules with the help of the CENP-E kinetochore motor (7-9).

The RZZ (Rod-ZW10-Zwilch) complex has been reported to be a key player in the spindle assembly checkpoint (SAC) activation as it is required to recruit SAC proteins, Mad1 and Mad2 to kinetochores in both *Drosophila* and humans (10-15). More importantly, it has also been shown that the RZZ complex is important to recruit dynein to kinetochores through its direct association with the dynein adaptor protein, Spindly (16-21). However, it is clear that there are also RZZ-independent mechanisms (such as the CENP-F-NudE pathway) contributing to this function (22,23). The dynein motor has been shown to be involved in rapid movement of mono-oriented chromosomes towards the spindle poles via dynamic lateral interaction between kinetochores and astral microtubules during early prometaphase, thus contributing to chromosome alignment (2,21,24-26). As these proteins are interlinked and function together at kinetochores in this process for maintaining dynamic kMT attachments, they constitute a module referred to as the “dynein module” (27).

The Ndc80 complex, consisting of four coiled-coil proteins Hec1, Nuf2, Spc24, and Spc25, is a major constituent of the outer plate of kinetochores, and is required for stable end-on kMT attachments after chromosome alignment at the metaphase plate (27-29). Recent studies in *C. elegans* have shown that the Rod subunit of the RZZ complex interacts with the Hec1 subunit of the Ndc80 complex and that this association is critical for forming stable kMT attachments during mitotic chromosome alignment. The presence of Rod at kinetochores was shown to be inhibitory for the formation of stable kMT attachments by the Ndc80 complex, possibly in the early stages of mitosis, to control the strength of kMT attachments in an Aurora B kinase-independent manner (17,27). The removal of SAC proteins, including Rod, from kinetochores by Spindly-dynein mediated “stripping” during checkpoint silencing is thought to enable Ndc80 to form stable kMT attachments at the spindle equator. In addition, super-resolution mapping of the kinetochore location of the components of the RZZ complex in humans suggests that they are located very proximal to the N-terminal region of the Ndc80 complex (15), which has been established to be critical for stable kMT attachment formation (28,30-34).

However, if/how the dynein module regulates kMT attachments of Ndc80 during early mitosis at human kinetochores to drive chromosome motility and alignment is unclear. Here, we address the functional relationship between the dynein module and the Ndc80 complex for chromosome alignment in human cells by using high-resolution confocal microscopy and siRNA-mediated functional perturbation studies. We find that the components of the dynein module serve key functions for regulating the stability of kMT attachments mediated by the Ndc80 complex.

## Results and Discussion

### Evidence for coordination between the dynein and Ndc80 kinetochore modules for proper chromosome alignment in humans

It is well-known that the attachments between kinetochores and kMTs in early mitosis are dynamic in nature to favor kinetochore MT-motor dependent chromosome motility that drives chromosome congression, and aid in attachment error correction (35). It is also established that the Ndc80 complex at kinetochores form strong attachments with spindle MTs to stabilize kMT attachments during chromosome alignment and bi-orientation at the spindle equator in metaphase (36,37), and that purified Ndc80 binds to microtubules with high affinity in vitro (30,38,39). Consistent with this, we find that relatively low concentrations of GFP-tagged Hec1/Nuf2 dimer of the Ndc80 complex (1-5 nM) bound readily to Dylight405-labeled MTs immobilized on coverslips in *in vitro* TIRF microscopy assays (Fig. 1*A*). In addition, it is known that the Ndc80 complex and the dynein motor share similar binding sites on MTs (40,41), which points to the notion that Ndc80-mediated kMT attachments may be mutually exclusive with dynein-based attachments during mitosis. We tested this directly by carrying out TIRF-M assays using GFP-tagged Hec1/Nuf2 dimer and TMR-labelled Dynein-Dynactin-BicD2 (DDB) (42). We found that the presence of higher concentrations of the Hec1/Nuf2 dimer (20 nM), strongly inhibited the binding of the DDB complex to MTs, supporting this prediction (Fig. 1, *B* and *C*). Consistent with this finding, it has been observed in *Xenopus* cells that the velocity of dynein-based poleward movement was substantially enhanced in the absence of the Ndc80 complex (26).

**FIGURE 1.**
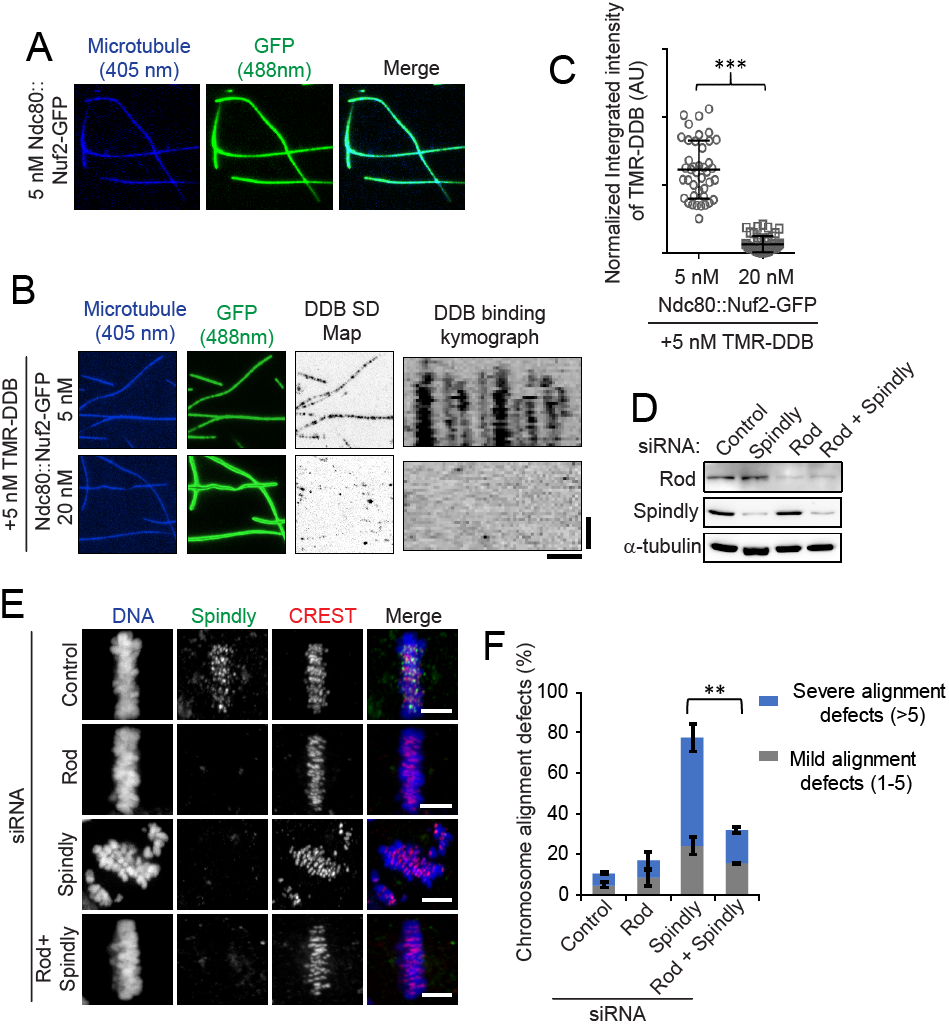
*A-C*, Higher conc of Ndc80 inhibits dynein MT-binding in in vitro TIRF microscopy assays. ***A***, Representative TIRF microscopy images showing surface-attached Dylight405-labeled MTs (blue), Ndc80::Nuf2-GFP (green) at 5nM. ***B***, Higher concentration of Ndc80::Nuf2-GFP (20 nM) (green) strongly inhibits the binding of TMR-DDB (red) on surface-attached Dylight405-labeled MTs (blue). The signal intensity of Ndc80::Nuf2-GFP is scaled identically between images. A standard deviation (SD) map of TMR-DDB binding over a 5 min movie is shown (59), along with a representative kymograph showing TMR-DDB binding on a single MT. In the absence of ATP, TMR-DDB binds statically to the MT lattice (vertical lines in kymograph). Bars, 5 μm, 2 min. ***C***, Quantification of integrated intensity of TMR-DDB on surface-attached MTs in the presence of variable concentration of Ndc80::Nuf2-GFP. Intensity was measured for at least 25 different locations along MTs for three independent images. ***P < 0.005 (Mann–Whitney U test. ***D-F*, Rescue of chromosome alignment defects of Spindly-depleted cells by codepletion of Rod. *D***, Western blot analysis of HeLa cells treated with siRNAs for control, Spindly, Rod, and Spindly + Rod. α-tubulin was used as a loading control. ***E***, Immunofluorescence staining of mitotic cells depleted of target proteins as indicated in comparison to control cells (top panel) for Spindly (green) and a kinetochore marker CREST (red) and counterstained with DAPI for DNA. Bars, 5 μm. ***F***, Quantification of mitotic cells with chromosome alignment defects in cells from **E**. Error bars represent S.D. from three independent experiments. For each experiment, 200 mitotic cells were examined. ***P < 0.005 (Student’s t test).

Our data supports the idea that the MT-binding activity of the Ndc80 complex could be directly affected by higher local concentrations of dynein at prometaphase kinetochores. We surmise that the initial capture and dynein-dependent poleward motility of kinetochores could thus be a natural bias for attachments that are dynamic in nature and a mechanism that prevents Ndc80-mediated stable attachments during early mitosis. While our data supports the hypothesis that dynein and Ndc80 directly affect each other’s MT-binding, it is also possible that there are mechanisms similar to that observed in *C. elegans* involving the components of the dynein module, including Spindly and Rod, that might modulate the function of Ndc80 in humans, and we sought to test this possibility next.

### Defects in chromosome alignment induced by Spindly or dynein depletion are rescued by codepletion of Rod

As Spindly has been shown to relieve the inhibition of Ndc80 by Rod to enable the formation of Ndc80-mediated stable kMT attachments in worms (17,27), we tested whether Rod is functionally related to Spindly in humans similar to that of the observation in worms. We sought to assess the phenotype of chromosome alignment in cells where the function of Rod and/or the dynein anchor, Spindly, was disrupted by RNAi-mediated knockdown of both the proteins. Efficient depletion of the target proteins was validated by immunoblotting as well as immunostaining analyses (Fig. 1, *D* and *E*; Supplemental Fig. S1*A*). We found that mitotic cells depleted of Rod (Rod^siRNA^) exhibit no apparent defect in chromosome alignment at the metaphase plate (Fig. 1*E*, Supplemental Fig. S1*A*). As observed in worms (17,27), the severe chromosome misalignment produced by Spindly depletion (Spindly^siRNA^) was rescued by the codepletion of Rod (Spindly/Rod^siRNA^) (Fig. 1, *E* and *F*; Supplemental Fig. S1*A*). The frequency of mitotic cells with misaligned chromosomes was significantly lower after Spindly/Rod^siRNA^, compared to that of Spindly^siRNA^ (Fig. 1*F*). These observations suggest that the modulation of Rod function by Spindly to aid in the formation of Ndc80-mediated stable kMT attachments is conserved from worms to humans; but the molecular mechanism of how this inhibition is accomplished is poorly understood.

Since dynein employs Spindly as an adaptor to bind to the RZZ complex and get recruited to kinetochores (17-19), we then tested if chromosome alignment defects after dynein^siRNA^ are also rescued by codepletion of Rod. To effectively deplete dynein, we designed a new siRNA targeting the 3’UTR sequence of the DYNC1H1 gene. Efficient depletion of the target proteins was validated by immunoblotting as well as immunostaining analyses (Fig. 2, *A-C*; Supplemental Fig. S1*A*). As expected (21,24,43), depletion of dynein caused a significant increase in the percentage of mitotic cells with misaligned chromosomes in HeLa cells (Fig. 2, *C* and *D*). On the other hand, the frequency of cells with misaligned chromosomes was significantly reduced after dynein/Rod^siRNA^ compared to that of dynein^siRNA^, and was almost similar to that of control^siRNA^ (Fig. 2, *C* and *D*; Supplemental Fig. S1*A*). As a positive control, depletion of Ndc80 using siRNA-mediated knockdown of the Hec1 subunit (Ndc80^siRNA^) expectedly led to severe chromosome alignment defects (44,45) (Fig. 2*D*; Supplemental Fig. S2, *A* and *B*). We further confirmed the rescue of chromosome alignment defects after dynein/Rod^siRNA^ by live imaging. We found that ∼80% of control cells could align their chromosomes at the metaphase plate within 30 min of the NEB (nuclear envelope breakdown) whereas ∼75 % dynein^siRNA^ cells were not able to do so even 120 min after the NEB. On the other hand, ∼60 % of dynein/Rod^siRNA^ cells could align their chromosomes with only a mild delay compared to that of control^siRNA^ cells (Fig. 2, *E* and *F*; Supplemental movies 1-4). As expected, we also observed severe chromosome alignment defects by live cell imaging after Ndc80^siRNA^ (Fig. 2*F*; Supplemental Fig. S2*C*; Supplemental movies 5 and 6). Thus, our results suggest that the defect in chromosome alignment resulting from dynein depletion was surprisingly rescued by codepletion of Rod.

**FIGURE 2.**
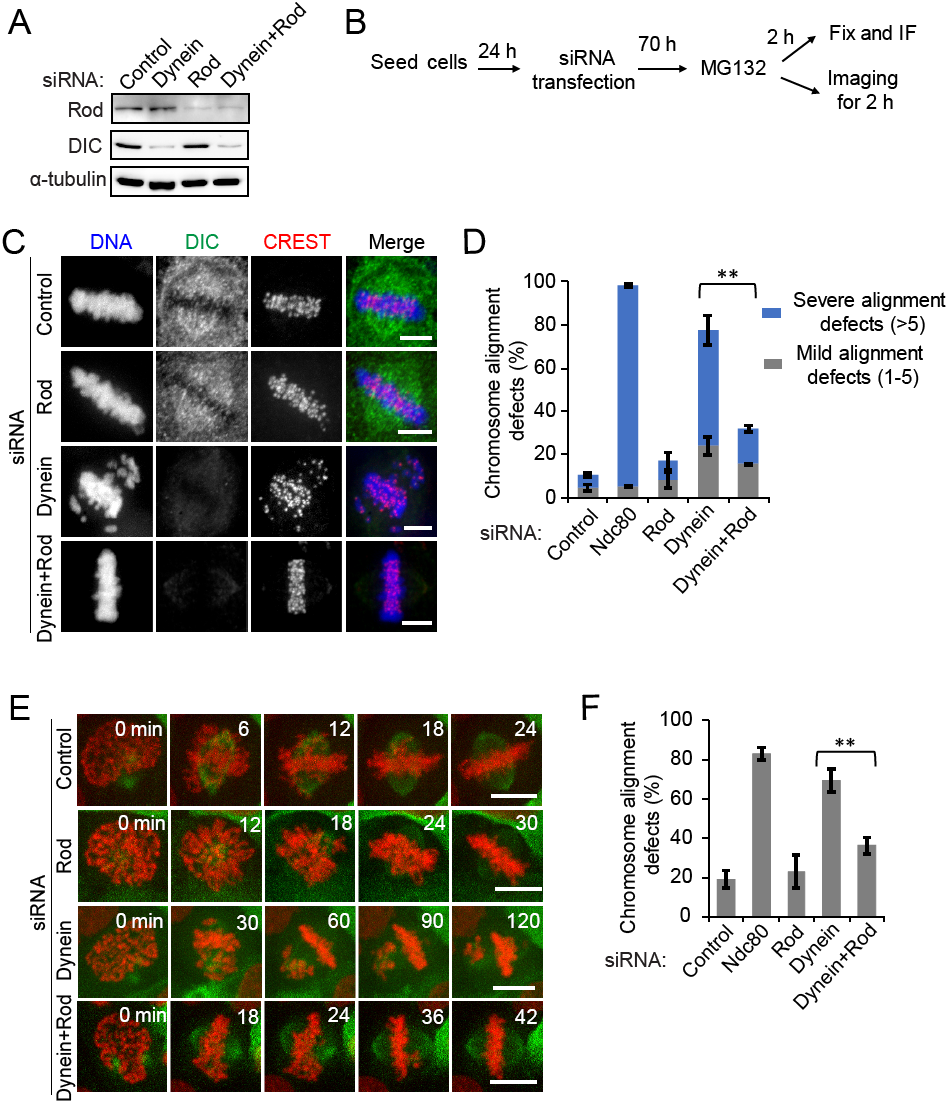
Rod codepletion rescues the defects in chromosome alignment induced by depletion of dynein. ***A***, Western blot analysis of HeLa cells treated with siRNAs for control, dynein, Rod and dynein + Rod. α-tubulin was used as a loading control. ***B***, Cells were transfected with the indicated siRNAs, and synchronized and fixed according to the scheme. ***C***, Immunofluorescence staining of mitotic cells depleted of target proteins as indicated in comparison to control cells (top panel) for dynein intermediate chain (DIC, green) and a kinetochore marker CREST (red) and counterstained with DAPI for DNA. Bars, 5 μm. ***D***, Quantification of mitotic cells with chromosome alignment defects in cells from ***C*** and in cells depleted of Ndc80 (also see Supplemental Fig. S2). Error bars represent S.D. from three independent experiments. For each experiment, 200 mitotic cells were examined. **P < 0.01 (Student’s t test). ***E***, Selected frames of videos from HeLa cells expressing H2B-mCherry and GFP-α-tubulin treated with siRNAs as indicated. Images were captured at 6 min intervals starting from nuclear envelope breakdown for 2 h, immediately after the addition of MG132. Scale bars: 10 μm. ***F***, Quantification of cells with chromosome alignment defects in ***E*** and of cells with chromosome alignment defects after Ndc80 depletion (also see Supplemental Fig. S2). Error bars represent S.D. from three independent experiments. For each experiment, 30 mitotic cells were examined.

The existing paradigm demonstrating a critical role for dynein in the rapid poleward movement of chromosomes during early mitosis to drive chromosome alignment originates from studies in large mitotic cells such as newt pneumocytes where the chromosomes are separated by relatively large distances (several 10s of μms) from the spindle poles (46). However, in smaller mitotic cells such as HeLa where the chromosomes are separated only by smaller distances for the spindle poles (usually within 5-10 μms), we surmise that the disengagement of dynein/spindly from Rod and Ndc80 is a major cause of chromosome misalignment after dynein‐ or spindly-depletion, due to which Rod is able to impart a sustained inhibition of Ndc80 function. It is also possible that the polar ejection forces on chromosome arms produced by the MT plus-end-directed chromokinesin motors drive the chromosomes away from the spindle poles during early mitosis in the absence of dynein or spindly to hinder proper chromosome alignment (4,47) (also see in Fig. 3).

**FIGURE 3.**
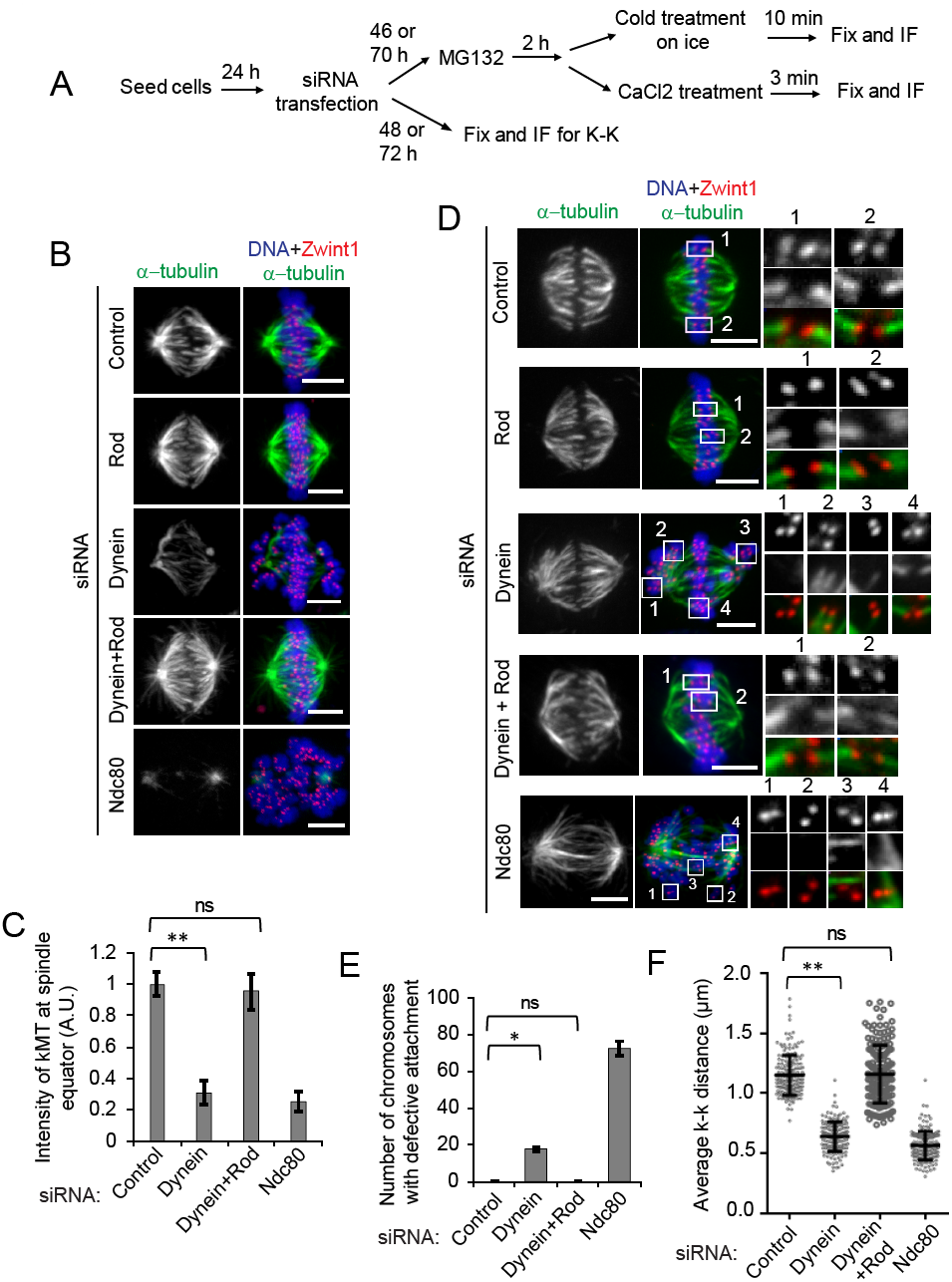
The formation of stable kMT kMT attachments in dynein-depleted cells is restored by codepletion of Rod. ***A***, Cells were transfected with the indicated siRNAs, synchronized, cold treated and fixed according to the scheme. ***B***, HeLa cells treated with siRNAs as indicated after cold treatment were immunostained for α-tubulin (green) and Zwint1 as a kinetochore marker (red), and counterstained with DAPI for DNA (blue). Bars, 5 μm. ***C***, Quantification of intensities of cold stable kMTs at the spindle equator of metaphase plate in ***B***. A.U.: arbitrary units. Error bars represent S.D. from three independent experiments. For each experiment, 10 cells were examined. ***P < 0.005 (Mann–Whitney U test), ns: not significant. ***D***, Analysis of end-on kMT attachments and kinetochore bi-orientation in cells treated with siRNAs as indicated. Cells were pre-treated with CaCl_2_ (0.2 mM) for 3 min to depolymerize unstable microtubules, and immunostained for α-tubulin (green) and Zwint1 as a kinetochore marker (red), and counterstained with DAPI for DNA (blue). Bars, 5 μm. ***E***, Quantification of the number of chromosomes with defective kMT attachments in ***D***. ***F***, Quantification of inter-kinetochore (k-k) distances measured from Zwint1 signals at sister kinetochore pairs of cells treated with siRNAs as indicated. Error bars represent S.D. from three independent experiments. For each experiment, 10 cells were examined. Distance was measured for at least 10 pairs of kinetochores from each cell. N = 150, **P < 0.01 (Mann–Whitney U test), ns: not significant.

### The restoration of proper chromosome alignment in Rod-depleted cells is independent of its function in the spindle assembly checkpoint activation

The RZZ complex is important to recruit the spindle assembly checkpoint (SAC) proteins, Mad1 and Mad2 to kinetochores in both *Drosophila* and humans (10-15). To scrutinize the possibility that the rescue of chromosome alignment after dynein/Rod^siRNA^ was not due to aberrant checkpoint silencing, we tested the recruitment of SAC protein Mad1 to kinetochores of mitotic cells in prometaphase. We found that detectable levels of Mad1 was still present at kinetochores in prometaphase cells depleted of Rod, dynein, or of both Rod and dynein (Supplemental Fig. S1*B*). When we analyzed mitotic progression by live imaging, we found no apparent defect in chromosome alignment at the metaphase plate after dynein/Rod^siRNA^ similar to that of Rod^siRNA^ (Fig. 2, *E* and *F*). Moreover, in both cases, live cells that were not treated with MG132 neither underwent premature anaphase onset during mitosis nor exhibited micro-nuclei formation in interphase (data not depicted), implying that the function of Rod in chromosome alignment could be independent of its role in SAC activation during mitosis in human cells. These data also suggest that dynein/Rod^siRNA^ cells can align their chromosomes properly without impairing checkpoint activation.

### The rescue of chromosome alignment defects after dynein and Rod codepletion is accompanied by restoration of attachment and stability of kMT

Proper chromosome alignment at the metaphase plate is accompanied by the stabilization of kMTs, which are MT bundles that extend from the bi-oriented kinetochores to spindle poles (48-50). Our immunostaining data showed that kMTs resistant to cold treatment were markedly reduced in mitotic cells after dynein^siRNA^ as compared to that of control^siRNA^ (Fig. 3, *A-C*). Surprisingly, we found that dynein/Rod^siRNA^ cells were able to form robust kMTs to a similar extent as was observed after control^siRNA^ or Rod^siRNA^ (Fig. 3, *B* and *C*). As a positive control, we observed a severe lack of cold-stable kMTs after Ndc80^siRNA^ (Fig. 3, *B* and *C*). These data suggest that Rod’s inhibition of Ndc80-mediated kMT attachments (27) is promoted in the absence of the counter-activity of dynein on Rod; as a consequence kMT attachments are unstable in dynein-depleted cells.

Analyses of kMT attachments in cells treated with 0.2mM CaCl_2_ followed by immunostaining showed that dynein^siRNA^ caused misaligned chromosomes with syntelic, monotelic and unattached kMT attachments, and reduced the average distance between sister kinetochore pairs (k-k distance) at the spindle equator of metaphase plate (Fig. 3, *D-F*). In the absence of the poleward-directed motor activity of dynein, we surmise that the misaligned chromosomes are driven to form syntelic and monotelic kMT attachments due to the activity of plus-end-directed chromokinesin motors (4,47). We found that the defects in kMT attachments and k-k distances resulting from dynein depletion were rescued by codepletion of Rod. The number of defective kMT attachments was significantly lower after dynein/Rod^siRNA^ compared to that of dynein^siRNA^, and was similar to that of control^siRNA^ (Fig. 3, *D* and *E*). The average k-k distances after dynein/Rod^siRNA^ was restored to that of control^siRNA^ (Fig. 3*F*), possibly due to the re-establishment of stable kMT attachments that we observed. As a positive control, Ndc80^siRNA^ cells exhibited severely defective kMT attachments and abnormal inter-kinetochore stretch (44,51) (Fig. 3, *D-E*). These results suggested that codepletion of Rod rescued the defects in chromosome alignment resulting from dynein depletion by restoring the robustness of kMTs, the stability of kMT attachments, and average interkinetochore distances.

**FIGURE 4.**
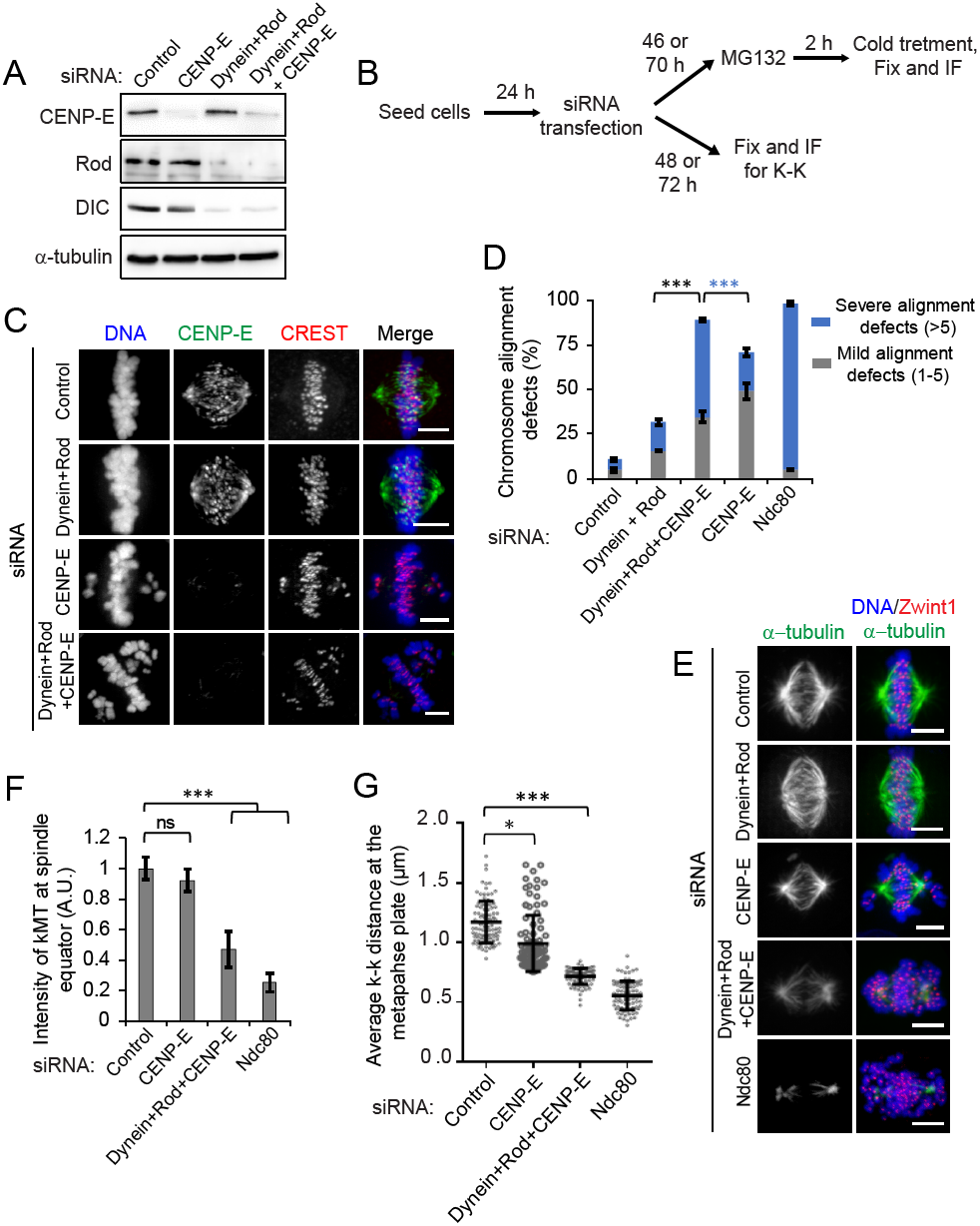
Chromosome alignment in cells codepleted of dynein and Rod is dependent on the motor activity of CENP-E. ***A***, Western blot analysis of HeLa cells treated with siRNAs for control, CENP-E, dynein + Rod and dynein + Rod + CENP-E. α-tubulin was used as a loading control. ***B***, Cells were transfected with the indicated siRNAs, synchronized, cold treated and fixed according to the scheme. ***C***, Immunofluorescence staining of mitotic cells depleted of target proteins as indicated in comparison to control cells (top panel) for CENP-E (green) and a kinetochore marker CREST (red) and counterstained with DAPI for DNA (blue). Bars, 5 μm. ***D***, Quantification of mitotic cells with chromosome alignment defects in cells from ***C*** and for cells treated with siRNA for Ndc80. Error bars represent S.D. from three independent experiments. For each experiment, 200 mitotic cells were examined. ***P < 0.005 (Student’s t test). ***E***, HeLa cells treated with ice-cold L-15 medium after treatment with siRNAs as indicated were immunostained for spindle MT (green) and a kinetochore marker, Zwint1 (red) and counterstained with DAPI for DNA (blue). Bars, 5 μm. ***F***, Quantification of intensities of kMTs adjacent to the kMT attachment sites. A.U.: arbitrary units. Error bars represent S.D. from the mean for three independent experiments. For each experiment, 10 cells were examined. **P < 0.01 and *P < 0.05 (Mann– Whitney U test). ***G***, Quantification of inter-kinetochore (k-k) distances measured from Zwint1 signals at kinetochore pairs of cells treated with siRNAs indicated. Error bars represent S.D. from three independent experiments. For each experiment, 10 cells were examined. Distance was measured for at least 10 pairs of kinetochores from each cell. n = 150, **P < 0.01 (Mann–Whitney U test), ns: not significant.

We hypothesize that during early mitosis, in the absence of the kinetochore dynein module, Ndc80 plays a role in the formation of initial kMT attachments, which are subsequently stabilized after chromosome alignment at the metaphase plate driven by CENP-E and/or chromokinesins. To support this prediction, we tested the status of kMT attachment of misaligned chromosomes in cells depleted of dynein or Ndc80. Close inspection of misaligned chromosomes showed that kinetochores were attached to MTs with syntelic or monotelic orientation after dynein^siRNA^ while those in Ndc80^siRNA^ cells remained unattached (Fig. 3*D*), suggesting that Ndc80 might have an unexplored, yet important role in initial kMT capture. These observations could also explain how chromosomes are still captured by spindle MTs after dynein/Rod^siRNA^. These results support the idea that a key function of Rod is to inhibit Ndc80 because kMT attachments are rescued when Rod is codepleted with either spindly (17,27,52) or dynein (this study).

Together, these data suggest that Rod is a negative regulator of stable kMT attachments and serves not only to perturb the function of the Ndc80 complex (27) but also that it’s inhibitory function is controlled by Spindly and dynein, the mechanism for which is yet unclear. The above data also suggest that chromosome alignment can be achieved at the spindle equator in the absence of the dynein module for human kinetochores, when normal chromosome alignment could possibly occur either by the activity of residual CENP-F/NudE-recruited dynein (22,23) and/or by the activity of the plus-end directed kinetochore motor, CENP-E.

### The chromosome alignment in cells codepleted of dynein and Rod is dependent on the motor activity of CENP-E

As normal chromosome alignment persists after dynein/Rod^siRNA^, we sought to investigate if the plus-end directed kinetochore motor CENP-E, which is involved in transporting unattached sister kinetochores along pre-existing microtubule bundles to the metaphase plate (4,8,53,54), was involved in chromosome congression by simultaneously perturbing CENP-E function using siRNA-mediated knockdown in dynein/Rod^siRNA^ cells. Efficient depletion of the target proteins was validated by immunoblotting as well as immunostaining analyses (Fig. 4, *A* and *C*; Supplemental Fig. S3, *A* and *B*). Immunostaining data showed that the frequency of cells with severe chromosome misalignment (more than 5 chromosomes) was significantly higher after dynein/Rod/CENP-E^siRNA^ (∼54%) as compared to that after control^siRNA^ (∼5%), dynein/Rod^siRNA^ (∼16%), or CENP-E^siRNA^ (∼ 21%) (Fig. 4, *C* and *D*; Supplemental Fig. S3, *A* and *B*). As a positive control described previously, ∼ 80% of cells after Ndc80^siRNA^ exhibited major chromosome misalignments (Supplemental Fig. S2, *A* and *B*). We further confirmed this result by inhibiting the motor activity of CENP-E using the small molecule inhibitor GSK-923295 (55), which we found, also abolished the rescue of chromosome alignment defects after dynein/Rod^siRNA^ (Supplemental Fig. S3, *C-E*).

The lack of rescue of chromosome alignment after dynein/Rod/CENP-E^siRNA^ prompted us to test whether stable kMT attachments were formed normally in these cells. We observed a substantial decrease in the intensity of kMTs at the spindle equator after dynein/Rod/CENP-E^siRNA^ similar to that of Ndc80^siRNA^ (Fig. 4, *E* and *F*), and in contrast to what was previously observed after dynein/Rod^siRNA^ or CENP-E^siRNA^ (Fig. 3, *B* and *C*; 4, *E* and *F*). We believe that the severe chromosome misalignment produced after dynein/Rod/CENP-E^siRNA^ prevents proper kinetochore biorientation, due to which the kMTs retain their cold sensitive nature. Moreover, the average inter-kinetochore distance was significantly reduced after dynein/Rod/CENP-E^siRNA^ in contrast to that of control^siRNA^ or CENP-E^siRNA^, and similar to that of Ndc80^siRNA^ (Fig. 4*G*). These observations suggest a biased mechanism for the rescue of chromosome alignment after dynein/Rod^siRNA^ that is mediated by the plus-end directed motility of CENP-E.

### Summary and conclusions

Overall, this study establishes a functional relationship between the kinetochore dynein module and Ndc80 module for proper chromosome alignment in humans. Rod is a key recruiter of the dynein module because it is involved in recruiting both dynein and Spindly to kinetochores (16,18,19,52,56). Our experimental results show that depletion of Rod alone has no apparent effect on chromosome alignment (Fig. 1, *E* and *F*). We found that severe defects of chromosome alignment after Spindly^siRNA^ were rescued by codepletion of Rod in humans similar to the observation in worms (17,27). Surprisingly, we also found that severe defects in chromosome alignment after dynein^siRNA^ were rescued by codepletion of Rod. As Spindly, a dynein adapter, counteracts the inhibitory role of Rod in the formation of Ndc80-mediated stable end-on kMT attachments (17,27), the analogous relationship between dynein and Rod; similar to that of Spindly and Rod, led us to propose that dynein could also directly counteract the inhibitory role of Rod on stable kMT attachments during early mitosis. Therefore, the phenotype of either Spindly^siRNA^ or dynein^siRNA^ represent the outcome of the disengagement of Rod from these factors, thus leading to severe chromosome alignment defects. Taken together these observations lead us to conclude that the role of dynein module in chromosome alignment depends on the function of Rod. It is not clear at this point whether it is the Rod/Spindly-dependent or CENP-F/NudE-dependent kinetochore dynein population, that is critical for the dynein-module mediated control of Ndc80 function in early mitosis.

Under normal condition after the NEB, kinetochores that are initially attached to spindle MTs by “search and capture” mechanism get rapidly transported poleward along MTs primarily by minus-end-directed motor force of dynein, and consequently chromosomes congress to the spindle equator by the activity of CENP-E motor (4). From recent studies (17,27 and this study), we propose a refined model for controlling kMT attachments during early mitosis in human cells, where Spindly inhibits Rod directly and/or through the dynein motor. Initial kMT attachments can still be formed by Ndc80 but are not stabilized due to antagonistic activity between the dynein module and Ndc80 (Fig. 5, details in legends). However, the precise mechanism for how Spindly/dynein interferes with Rod function and how Rod inhibits Ndc80 is stillunclear. The initial kMT attachments allow dynein-mediated transport of chromosomes to the spindle pole in early mitosis during which both CENP-E activity and end-on attachment formation are expected not to be favored (7). Therefore, in cells codepleted of dynein and Rod during early mitosis, initial kMT attachments can be formed by Ndc80 and chromosomes can be transported poleward by shrinkage of peripheral MT bundles, after which they get congressed at the metaphase plate mediated by CENP-E activity. Our current data leads us to propose the existence of two possibly interconnected mechanisms to control the stability of kMT attachments by the Ndc80 complex. Dynein might inhibit Rod to aid in Ndc80-mediated initial kMT attachments, while at the same time, or in an independent manner, directly inhibit the binding of the Ndc80 complex to MTs to prevent premature kMT attachment stabilization during in early mitosis.

**FIGURE 5.**
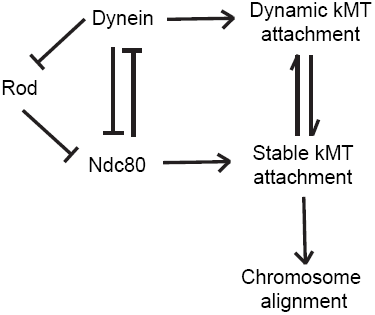
A model integrating the mechanisms that control stable kMT attachments in humans. During early mitosis, Rod, a subunit of the RZZ complex, which is involved in the recruitment of both dynein and its adaptor, Spindly, inhibits Ndc80 to prevent the formation of premature stable kMT attachments. Dynein and Spindly, which are involved in dynamic kMT attachments, inhibit Rod by a yet unclear mechanism, to modulate Rod-mediated inhibition of Ndc80 in stabilizing kMT attachments. Ndc80 is able to directly inhibit dynein binding to microtubules (MTs) while higher levels of dynein at prometaphase kinetochores might reciprocally also interfere with Ndc80-MT binding. Thus dynein-mediated dynamic kMT attachments and Ndc80-mediated stable kMT attachments are coordinated during early mitosis either by a direct interplay between dynein and Ndc80 and/or by the mediation of Rod to drive proper chromosome alignment.

Thus, it is becoming clear that stable kMT attachments are regulated not only by components within the dynein module but also by a direct interplay between the dynein module and the Ndc80 complex to prevent premature kMT stabilization. Thus, our results reveal a further layer of the elaborate network involving these distinct modules that are under tight spatiotemporal regulation to control kMT attachments and drive proper chromosome alignment at the metaphase plate. It will be important to continue studying the relationship between these modules with each other and with the protein complexes at the plus-ends of MTs in an effort to understand how they might coordinate the events involved in the formation and maturation of kMT attachments during chromosome alignment and segregation.

## Materials and methods

### Cell culture, transfections and drug treatments

HeLa cells were grown in Dulbecco’s modified Eagle medium (DMEM, Life Technologies) supplemented with 10% fetal bovine serum at 37 °C in humidified atmosphere with 5% CO_2_. For RNA interference (RNAi) experiments, cells were transfected at 30-50% confluence using Dharmafect 2 (Dharmacon) according to the manufacturer’s instructions, and analyzed 48-72 h after transfection. For rescue experiments, RNAi-refractory constructs were transfected into cells with Lipofectamine 3000 (Life Technologies) for 12 h followed by siRNA transfection. Cells were prepared for analysis after 48 h of siRNA transfection.

To arrest at metaphase, cells were treated with MG132 (10 μM) respectively for 2 h prior to fixation. To inhibit CENP-E, 200 nM GSK-923295 (APExBio) was added to the medium and cells were fixed after 2 h of incubation.

siRNAs targeting cDNAs for Ndc80 5’-GUUCAAAAGCUGGAUGAUCdTdT-3’ (45), Rod Invitrogen Stealth (5’-GGAAUGAUAUUGAGCUGCUAACAAA-3’) (57) and CENP-E Invitrogen stealth 5’-UUAUAUUACAGCCUUCCUGAGCCG-3’ (58) were used this study. A siRNA sequence targeting 3’ UTR for Dynein (5’-GGUGGAAAUUGGAAGGAUAdTdT-3’) was also used (Invitrogen). The Smart Pool ON-TARGET plus siRNAs used for targeting Spindly was purchased from Dharmacon (5’-GGGAGAAGUUUAUCGAUUA-3’, 5’-GAAAGGGUCUCAAA CUGAA-3’, 5’-GGAUAAAUGUCGUAAUGAA-3’, and 5’-CAGGUUAGCUGUUGAAUCA-3’). All siRNAs were used at 100 nM concentration except for dynein (200 nM).

### Antibodies

The primary monoclonal mouse antibodies used were: anti-Hec1 9G3 (Abcam, ab3613) for IF at 1:400, for WB at 1:1000; anti-Dynein clone 74.1 (EMD Millipore, MAB1618) for IF at 1:300, for WB at 1:1000, anti-Rod clone 10H4 (Abnova, H00009735-M01) IF at 1:300, WB at 1:1000; anti-CENP-E 1H12 (Abcam, ab5093) IF at 1:400, and anti-α-tubulin DM1A (Santa Cruz, Sc32293) IF at 1:750, WB at 1:1000. The primary rabbit polyclonal antibodies used were: anti-α-tubulin (Abcam, ab18251) for IF at 1:750; anti-Zwint1 (Bethyl, A300-781A) for IF at 1:400; anti-Spindly (a gift from Dr. Arshad Desai, UC San Diego) for IF at 1:5000; anti-Spindly (Bethyl, A301-354AT) for WB at 1:1000; anti-CENP-E (Bethyl, A301-943A) for WB at 1:1000. We also used a human anti-centromeric (Immunovision) at 1:1000 for IF.

### Immunofluorescence

HeLa cells grown on a glass coverslip were fixed in cold methanol (-20^o^C) or 4% formaldehyde after pre-extraction with 0.1% Triton X-100 followed by blocking with 3% bovine serum albumin (BSA) in PBS, and incubated with primary antibodies for 1 h at 37 ^o^C followed by washing with PBS (137mM NaCl, 2.7mM KCl, 10mM Na_2_HPO_4_ and 1.8mM KH_2_PO_4_, pH 7.4) supplemented with 0.02% Triton X-100. The primary antibodies were detected by using secondary antibodies coupled with Alexa-Fluor-488/647 or Rhodamine Red-X (Jackson ImmunoResearch Laboratories, Inc.) at a dilution of 1:250, and DNA was counterstained with 1 mg/ml DAPI.

For the analysis of cold-stable microtubules, cells were incubated for 10 min on ice in Leibovitz’s L-15 medium (Gibco) supplemented with 20 mM HEPES, pH 7.0 and 10% FBS, and then fixed for 15 min with 4% formaldehyde in PBS at 37 ^o^C. For the analysis of CaCl_2_-resistant microtubules, cells were treated with PCM buffer (60 mM PIPES, 0.2 mM CaCl_2_ and 4 mM MgSO_4_, pH 6.9) for 3 min and then fixed for 15 min at room temperature with 4% formaldehyde and 0.5% Triton X-100 in PCM buffer

### Image acquisition and analysis

For image acquisition, three-dimensional stacks were obtained through the cell using a Nikon Eclipse TiE inverted microscope equipped with a Yokogawa CSU-X1 spinning disc, an Andor iXon Ultra888 EMCCD camera and an x60 or x100 1.4 NA Plan-Apochromatic DIC oil immersion objective (Nikon). For fixed cell experiments, images were acquired at room temperature as Z-stacks at 0.2 μm intervals controlled by NIS-elements software (Nikon). Images were processed in Fiji ImageJ and Adobe Photoshop CC 2015 and represent maximum-intensity projections of the required z-stacks. To measure the fluorescence intensity, we selected manually and measured individual kinetochores by quantification of the pixel gray levels of the focused z plane in a region of interest (ROI) using Fiji ImageJ. The final intensity was obtained after subtracting the background, which was measured outside of the ROI, from the integrated intensity for the experiment. For the measurement of K-fiber intensity, we converted identically scaled images into TIFF files, selected ROIs at the equatorial position of the metaphase plate and measured microtubule intensity as described above.

### Live-cell imaging

HeLa cells stably expressing mCherry H2B and GFP-α-tubulin or only GFP-H2B were cultured in 35mm glass-bottomed dishes (MatTek Corporation). Before 30 min of imaging, cell culture medium was changed to pre-warmed L-15 medium (Gibco) supplemented with 20% fetal bovine serum and 20mM HEPES, pH 7.0. Live-experiments were carried-out in an incubation chamber for microscopes (Tokai Hit Co., Ltd) at 37 ^o^C and 5% CO_2_. Images recording was started immediately after adding MG132 (unless otherwise stated) using an x60 1.4 NA Plan-Apochromatic DIC oil immersion objective mounted on an inverted microscope (Nikon) equipped with an Andor iXon Ultra888 EMCCD camera or an Andor Zyla 4.2 plus sCMOS camera. Twelve 1.2 μm-separated z-planes covering the entire volume of the cell were collected at every 10 min up to 12 h. Images were processed in Fiji ImageJ and Adobe Photoshop CC 2015 and represent maximum-intensity projections of the required z-stacks.

### TIRF microscopy

All imaging experiments were performed at room temperature using a Nikon Eclipse Ti-E microscope equipped with a x100 1.49 numerical aperture oil-immersion objective and 1.5x tube lens (yielding a pixel size of 106 nm); an Andor iXon EM CCD camera; four laser lines (405, 491, 568 and 647 nm); and using the Micro-Manager 1.4.22 software. Flow chambers containing immobilized microtubules were assembled as described previously (42). All assays were performed in an assay buffer (90 mM HEPES of pH 7.4, 50 mM K-acetate, 2 mM Mg-acetate, 1 mM EGTA, and 10% glycerol) supplemented with 0.1 mg/mL biotin-BSA, 0.5% Pluronic F-168, and 0.2 mg/mL κ-casein. TMR-labeled DDB and Ndc80::Nuf2-GFP proteins were mixed together and flowed into the chamber and binding events were imaged over a period of 5 min. For the measurement of DDB intensity on the MT lattice, we converted identically scaled images into TIFF files, selected ROI on the MT lattice and obtained final DDB intensity by subtracting the background intensity measured outside of ROI from the integrated of DDB using FIJI/ImageJ. To generate a standard-deviation (SD) map of DDB binding, the SD projection option was selected within the Z-project tool in FIJI (59).

### Immunoprecipitation and immunoblotting

Cell lysates were prepared with lysis buffer [150 mM KCl, 75 mM HEPES of pH 7.5, 1.5 mM EGTA, 1.5 mM MgCl_2_, 10% glycerol, 0.1% NP-40, 30 mg/ml DNase, 30 mg/ml RNase, complete protease inhibitor cocktail (Roche) and complete phosphatase inhibitor cocktail (Sigma)]. Protein concentration of cell lysate was measured using the Coomassie protein assay kit (Thermo Scientific). For immunoprecipitation assays, pre-cleared native protein extracts (1 mg of total protein) were incubated overnight with mouse anti-Dynein clone 74.1 (EMD Millipore, MAB1618) at a dilution of 1:100 or mouse nonspecific IgG at a dilution of 1:100 (Invitrogen), followed by incubation with 40 μl of anti-mouse-IgG magnetic Dynabeads (Invitrogen) for 2 hours at 4°C. After washing three times with lysis buffer and once with cold 1xTBST, precipitated proteins were eluted by boiling for 5 minutes in 6xSDS sample buffer. Proteins were separated on SDS-PAGE, electroblotted onto a nitrocellulose blotting membrane (Amersham, GE Healthcare) and subjected to immunodetection using appropriate primary antibodies. Blocking and antibody incubations were performed in 5% non-fat dry milk. Proteins were visualized using horseradish peroxidase-conjugated secondary antibodies diluted at 1:2,000 (Amersham) and the ECL system, according to the manufacturer’s instructions (Thermo Scientific).

### Statistical analysis

Mann–Whitney U-test was used for comparison of dispersion, and a two-sided t-test was used for comparison of average. The statistical analyses were done with Prism software (GraphPad). Samples for analysis in each data set were acquired in the same experiment, and all samples were calculated at the same time for each data set.

## Acknowledgements

The Authors would like to thank Dhanya Cheerambathur and Arshad Desai for providing Ndc80 (Hec1)::Nuf2-GFP dimeric proteins of the Ndc80 complex This work was supported by an NCI grant to DV (R00CA178188) and by start-up funds from Northwestern University.

## Conflict of Interest

The authors declare no competing financial interest.

## Abbreviations List

NEB, nuclear envelope breakdown; DIC, Dynein intermediate chain, kMT, kinetochore microtubule; RZZ, Rod-Zw10-Zwilch; SAC, spindle assembly checkpoint; Ndc80, nuclear division cycle 80; CENP-E, Centrosome-associated protein E; ACA (CREST), anti-centromere antiserum; siRNA, small interfering RNA; KMN, Knl1-Mis12-Ndc80

## Author’s Contributions

M.A. and D.V. designed and conceived the research. M.A performed all the experiments and data analysis, except that R.J.M. performed the experiments and analysis involving TIRF microscopy. M.A. and D.V. contributed to writing the manuscript. M.A, D.V., and R.J.M contributed to editing the manuscript.

## Supplemental Figure Legends

**SUPPLEMENTAL FIGURE S1.**
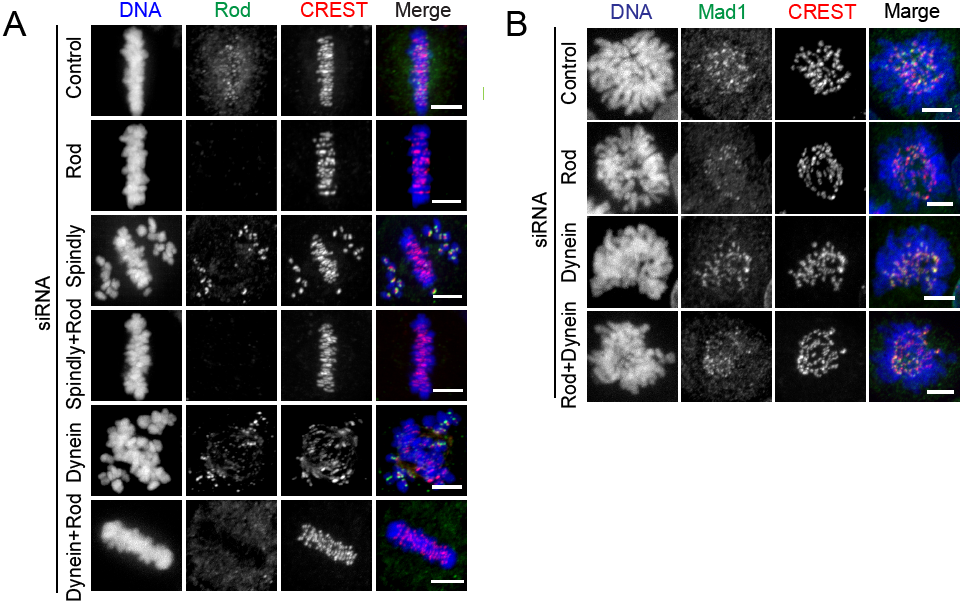
*A*, Phenotypic characterization of Dynein, Spindly or Dynein/Spindly and Rod double-depleted cells. HeLa cells treated with siRNAs as indicated were immunostained for α-tubulin (green) and a kinetochore marker CREST (red), and counterstained with DAPI for DNA (blue). Cells were treated with 10 μM MG132 for 2 h. Scale bars: 5 μm. ***B***, **Analysis of checkpoint protein activation.** HeLa cells treated with siRNAs as indicated were immunostained for α-tubulin (green) and a kinetochore marker CREST (red), and counterstained with DAPI for DNA (blue). Bars: 5 μm.

**SUPPLEMENTAL FIGURE S2.**
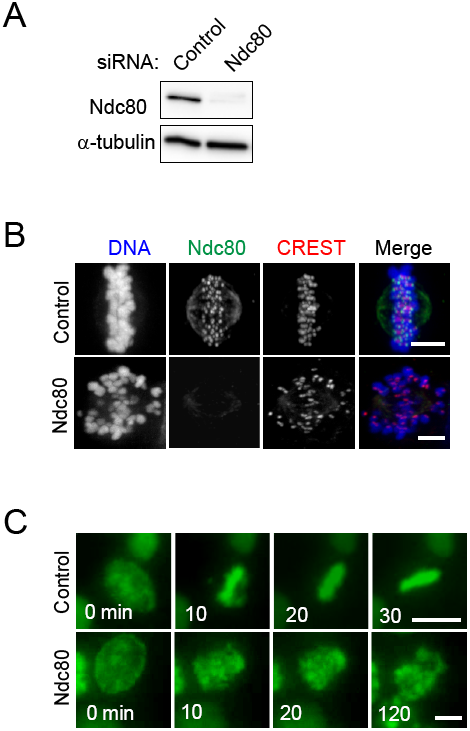
Chromosome alignment defects in cells depleted of Ndc80. ***A***, Western blot analysis of HeLa cells treated with siRNAs for control and Ndc80. α-tubulin was used as a loading control. ***B***, Immunofluorescence staining of mitotic cells depleted of Ndc80 in comparison to control cells (top panel) for Ndc80 (green) and a kinetochore marker CREST (red) and counterstained with DAPI for DNA. Bars, 5 μm. ***C***, Still images from live imaging of HeLa cells stably expressing histone H2B-GFP to visualize the chromosomes after control (top panel) or Ndc80 siRNA treatment (bottom panel). Imaging was started immediately after adding MG132 and images were captured at every 10 min for 2 h. Bars, 10 μm.

**SUPPLEMENTAL FIGURE S3.**
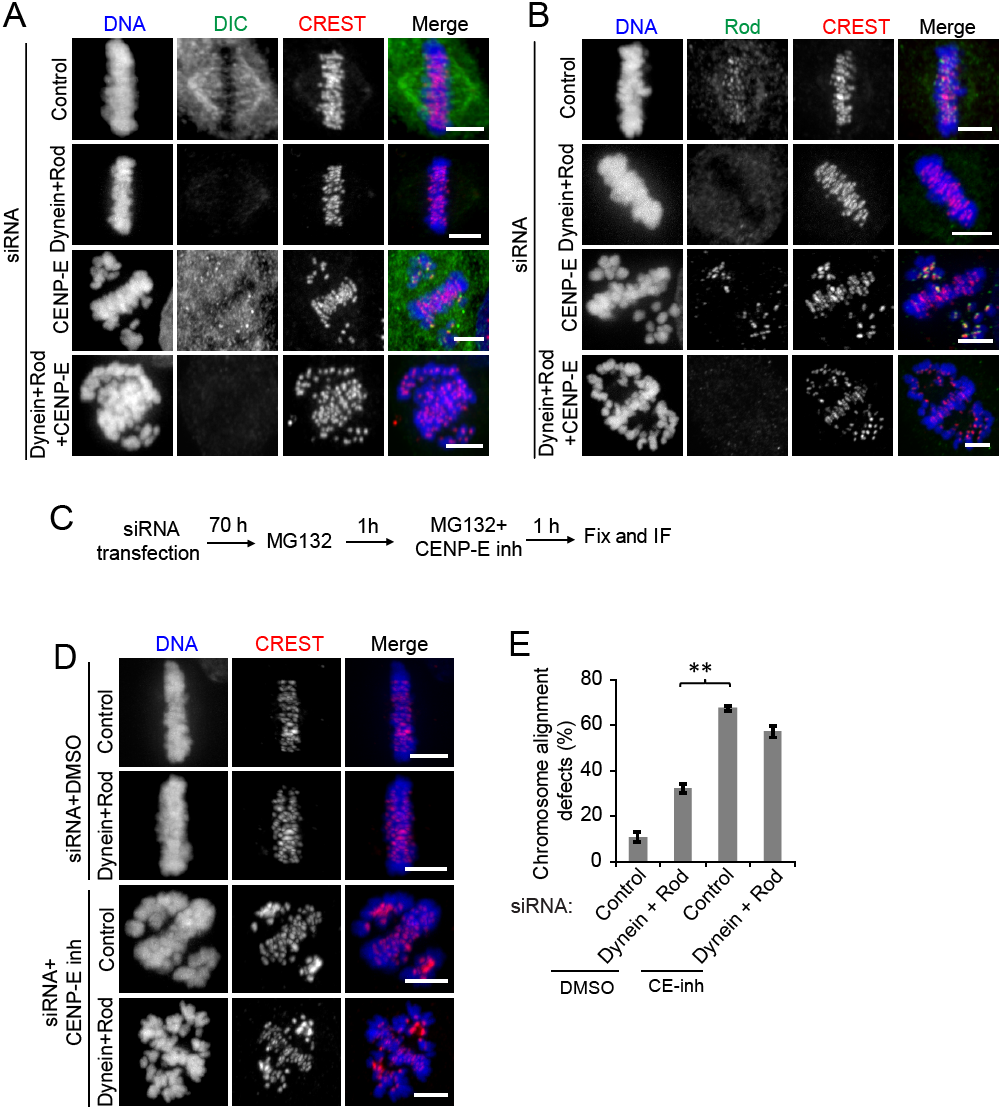
Phenotypic characterization of CENP-E depletion in the background of dynein and Rod double depletion. ***A*** and ***B***, HeLa cells treated with siRNAs as indicated were immunostained for DIC (***A***) and Rod (***B***) in green, a kinetochore marker CREST (red), and counterstained with DAPI for DNA (blue). Cells were treated with 10 μM MG132 for 2 h. Bars, 5 μm. ***C***, Cells were transfected with the indicated siRNAs, synchronized, and fixed according to the scheme. ***D***, HeLa cells treated with siRNAs as indicated were immunostained for a kinetochore marker CREST (red), and counterstained with DAPI for DNA (blue). Bars, 5 μm. ***E***, Quantification of chromosome alignment defects in cells from ***D***.

Supplemental Movie 1. Video for the chromosome alignment of HeLa cells expressing H2B-mCherry and GFP-α-tubulin treated with control siRNA. Imaging was started immediately after adding MG132 and continued for 2 h. Images were captured on a Nikon Eclipse TiE inverted microscope as z stacks of 1.2 μm thickness at 6 min interval starting from the nuclear envelope breakdown. Bars, 10 μm. Movie speed is 10 frames per second.

Supplemental Movie 2. Video for the chromosome alignment of HeLa cells expressing H2B-mCherry and GFP-α-tubulin treated with siRNA for dynein. Imaging was started immediately after adding MG132 and continued for 2 h. Images were captured on a Nikon Eclipse TiE inverted microscope as z stacks of 1.2 μm thickness at 6 min interval starting from the nuclear envelope breakdown. Bars, 10 μm. Movie speed is 10 frames per second.

Supplemental Movie 3. Video for the chromosome alignment of HeLa cells expressing H2B-mCherry and GFP-α-tubulin treated with siRNA for Rod. Imaging was started immediately after adding MG132 and continued for 2 h. Images were captured on a Nikon Eclipse TiE inverted microscope as z stacks of 1.2 μm thickness at 6 min interval starting from the nuclear envelope breakdown. Bars, 10 μm. Movie speed is 10 frames per second.

Supplemental Movie 4. Video for the chromosome alignment of HeLa cells expressing H2B-mCherry and GFP-α-tubulin treated with siRNA for dynein and Rod. Imaging was started immediately after adding MG132 and continued for 2 h. Images were captured on a Nikon Eclipse TiE inverted microscope as z stacks of 1.2 μm thickness at 6 min interval starting from the nuclear envelope breakdown. Bars, 10 μm. Movie speed is 10 frames per second.

Supplemental Movie 5. Mitotic progression in HeLa cells expressing Histone H2B-GFP treated with control siRNA. Imaging was started immediately after adding MG132 and continued for 2 h. Images were acquired on a Nikon Eclipse TiE inverted microscope as z stacks of 1.2 μm thickness every 10 minutes. Bars, 5 μm. Movie speed is 10 frames per second.

Supplemental Movie 6. Mitotic progression in Rod-depleted HeLa cells expressing Histone H2BGFP treated with siRNA for Ndc80. Imaging was started immediately after adding MG132 and continued for 2 h. Images were acquired on a Nikon Eclipse TiE inverted microscope as z stacks of 1.2 μm thickness every 10 minutes. Bar, 5 μm. Movie speed is 10 frames per second.

